# A Model Mechanism Based Explanation of an In Vitro-In Vivo Disconnect for Improving Extrapolation and Translation

**DOI:** 10.1101/216556

**Authors:** Andrew K. Smith, Yanli Xu, Glen E.P. Ropella, C. Anthony Hunt

**Affiliations:** Bioengineering and Therapeutic Sciences, University of California, San Francisco, CA 94143 (A.K.S., Y.X., C.A.H.); Tempus Dictum, Inc., Milwaukie, OR 97222 (G.E.P.R)

## Abstract

An improved understanding of in vivo-to-in vitro hepatocyte changes is crucial to interpreting in vitro data correctly and further improving hepatocyte-based in vitro-to-in vivo extrapolations to human targets. We demonstrate using virtual experiments as a means to help untangle plausible causes of inaccurate extrapolations. We start with virtual mice that have biomimetic software livers. Earlier, using those mice, we discovered model mechanisms that enabled achieving quantitative validation targets while also providing plausible causal explanations for temporal characteristics of acetaminophen hepatotoxicity. We isolated virtual hepatocytes, created a virtual culture, and then conducted dose-response experiments in both culture and mice. We expected the two dose-response curves to be displaced. We were surprised that they crossed because it evidenced that simulated acetaminophen metabolism and toxicity are different for virtual culture and mouse contexts even though individual hepatocyte mechanisms were unchanged. Crossing dose-response curves is a virtual example of an in vivo-to-in vitro disconnect. We use detailed results of experiments to explain the disconnect. Individual hepatocytes contribute differently to system level phenomena. In liver, hepatocytes are exposed to acetaminophen sequentially. Relative production of the reactive acetaminophen metabolite is largest (smallest) in pericentral (periportal) hepatocytes. Because that sequential exposure is absent in culture, hepatocytes from different lobular locations do not respond the same. A virtual Culture-to-Mouse translation can stand as a scientifically challengeable theory explaining an in vitro-in vivo disconnect. It provides a framework to develop more reliable interpretations of in vitro observations, which then may be used to improve extrapolations.

**Abbreviations:**

aHPC: analog hepatocyte
APAP: acetaminophen
CV: Central Vein
SS: sinusoidal segment
NAPQI: N-acetyl-p-benzoquinone imine
mitoD: mitochondrial damage products
nonMD: non-mitochondrial damage products

## Introduction

Results of quantitative hepatocyte-based in vitro-to-in vivo extrapolations continue to improve [Tetsuka et al., 2017, Poulin et al., 2016]. In vitro strategies for liver toxicity testing have experienced concurrent advances [Soldatow et al., 2013] aided by increasing knowledge about factors that limit accuracy [Vellonen et al., 2014, Fraczek et al., 2013, Godoy et al., 2013, LeCluyse et al., 2012]. Nevertheless, even for straightforward predictions of hepatic clearance from in vitro intrinsic clearance values, published results are typically more than 40% outside of in vivo values [Bowman and Benet, 2016], and some are 2-fold or more.

Different factors can contribute to inaccurate extrapolations. The following are five examples: 1) variability among livers; 2) periportal-to-pericentral hepatocyte differences; 3) variation in relative numbers of periportal, midzonal, and pericentral hepatocytes in cultured populations; 4) up- and down-regulation of genes during and after isolation; and 5) variation in hepatocyte health and phenotype caused by isolation and/or culture protocol differences. The prospect of being able to limit or avoid such changes is motivating interest in 3D hepatocyte cultures and microfluidic in vitro systems, with the expectation that they can become reliably more predictive of in vivo and human hepatic phenotypes.

An improved understanding of in vivo-to-in vitro hepatocyte changes is crucial to interpreting in vitro data correctly. We conjecture that insights gained from virtual experiments can help untangle and identify causes of inaccurate extrapolations to in vivo and human targets. The resulting new knowledge can then be used to improve in vitro-to-in vivo extrapolation methods. Figure 1 illustrates the key ideas. There are four essential requisites. 1) One can create many concrete, model mechanism-based, individualizable virtual hepatocytes that are demonstrably analogous to actual hepatocytes in particular ways. 2) The 3D organization of virtual hepatocytes is analogous to a rodent and human liver. 3) We achieve specific qualitative and quantitative validation targets for virtual counterparts of xenobiotics by changing the configurations of virtual hepatic mechanisms. 4) It is straightforward to “isolate” *all* virtual hepatocytes from the liver, without altering intra-hepatocyte mechanisms; and then reorganize them to simulate a 2D hepatocyte culture to study responses to virtual counterparts of xenobiotics. We recently reported using experiments on virtual mice to discover model mechanisms that together provide plausible causal explanations for major temporal characteristics of acetaminophen (APAP) hepatotoxicity in mice [Smith et al., 2016]. Those virtual mice meet the first three requisites. They utilize a software liver analog that is biomimetic across relevant anatomical, hepatic zonation, and cell biology attributes. Quasi-autonomous virtual hepatocytes populate virtual liver lobules. Each virtual hepatocyte uses the local values of periportal-to-pericentral gradients to configure its mechanisms for reactive metabolite formation, Glutathione (GSH) depletion, accumulation of mitochondrial damage, and triggering necrosis.

**Figure. 1.**
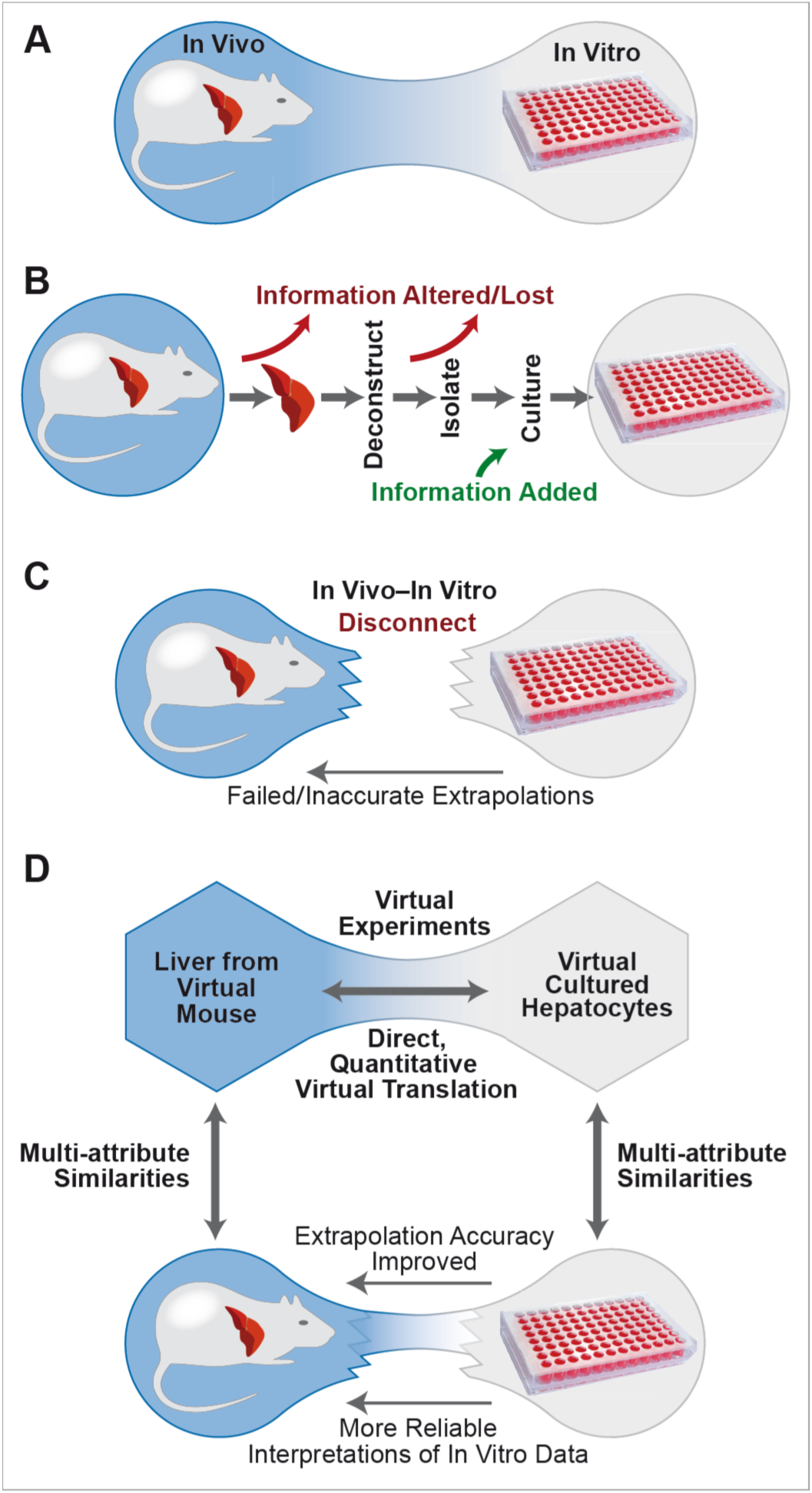
An approach to improve in vitro-to-in vivo extrapolations: the focus is mechanism-based explanations for differences in phenomena between two model systems. (A) An illustration of an ideal relationship: all phenotypic differences between in vivo and in vitro are understood; therefore, predictions and translations are knowledge based. (B) Some of the knowledge and information required to explain hepatocyte attributes (e.g., metabolic clearance) in vivo is lost and/or added during hepatocyte isolation and culture. (C) The processes illustrated in B alter hepatocyte phenotype, creating an in vivo-in vitro disconnect. An extrapolation or prediction based on an altered phenotype is inaccurate or fails to meet requirements. (D) Arrows on each side indicate multi-attribute similarities: each virtual system independently builds credibility through cumulative achievement of qualitative and quantitative referent system validation targets. Top – Differences in phenomena measured during Virtual Hepatocyte Culture experiments and corresponding measurements of phenomena recorded during Virtual Mouse experiments are entirely explainable: there is no in vivo-in vitro disconnect. Bottom – Mechanism differences enabling the virtual translation in the top illustration can stand as a concrete, scientifically challengeable theory explaining the disconnect in C. That model mechanism-based theory provides the framework to develop more reliable interpretations of in vitro observations, which then may be used to improve extrapolations.

An objective of this research was to demonstrate the feasibility of achieving requisite (4). An early, working hypothesis was that simulated APAP metabolism and toxicity would be essentially the same for virtual culture and mouse experiments.

We created a virtual hepatocyte culture and verified that it met requisite (4). We conducted dose-response experiments to challenge our working hypothesis. Because simulated transformation to culture configuration eliminates lobular structures, we expected dose-response curves to be somewhat displaced from each other but have similar shapes. We were initially surprised that the two curves crossed because it evidenced that simulated APAP metabolism and toxicity are different for virtual culture and mouse contexts even though individual hepatocyte mechanisms were unchanged. That evidence falsified our working hypothesis. Crossing dose-response curves is a virtual example of an in vivo-to-in vitro disconnect (Fig. 1C).

We use results of experiments to explain the apparent disconnect in detail (Fig. 1D). Individual hepatocytes exposed to the same amounts of APAP respond the same in both contexts. However, the relative contribution of different individual hepatocytes to the system level phenomenon is different. In liver, hepatocytes are exposed to APAP sequentially, periportal to pericentral, and both APAP intrinsic clearance and relative production of reactive metabolite are largest (smallest) in pericentral (periportal) hepatocytes. That sequential exposure is absent in culture. An important characteristic of sequential exposure is that relative numbers of hepatocytes are periportal > midzonal > pericentral. Accordingly, hepatocytes from different lobular locations do not respond the same when studied in culture.

We suggest that virtual translational experiments, like those described herein, can be used to begin exploring, untangling, and identifying causes of inaccurate extrapolations. Those improved insights can guide selecting additional in vitro measurements to enable more reliable interpretations of in vitro data. The approach provides a new, knowledge- and mechanism-grounded means to begin closing in vitro-in vivo disconnects.

## Material and Methods

Our virtual mouse is engineered to have software components that are concrete and strongly analogous to counterpart mouse components, but only to the extent needed to achieve prespecified by Targeted Attributes [Smith et al., 2016]. To stress that analogies—although numerous, qualitative, and quantitative—are limited, we refer to the virtual mouse as Mouse Analog. To limit confusion hereafter and distinguish Mouse (Culture) Analog components, characteristics, and phenomena from mouse (culture) counterparts, we capitalized the former. We used the scientific method to falsify three plausible model mechanisms for APAP Hepatotoxicity and discover a fourth, which is the one used here. To support the model mechanism-based explanations of results that follow, we provide abridged descriptions of Mouse Analog, intra-Hepatocyte Mechanisms, requirements, and technical details.

### Mouse Analog

Mouse Analog, which is illustrated in Fig. 2A, comprises Liver, Body, and a space to contain Dose. Liver, which uses Monte Carlo-determined Lobule variants, is designed and built to be scientifically useful in a variety of usage contexts, including developing model mechanism explanations of drug-induced liver injury. It is engineered to be quantitatively and qualitatively biomimetic during execution and is strongly analogous to actual livers across several anatomical, hepatic zonation, and cell biology characteristics. Having already achieved several qualitative and quantitative Target Attributes [Smith et al., 2016, Smith et al., 2014, Yan et al., 2008a, Yan et al., 2008b, Park et al., 2009, Park et al., 2010], Liver composition is now stable and robust. In this work, a Target Attribute is a characteristic trait of the liver and APAP metabolism, disposition, and toxicity to which a prespecified Similarity Criterion is assigned. The latter is a performance requirement. Each wet-lab measurement that we seek to mimic, such as hepatic extraction ratio, clearance, metabolite ratios, and necrosis, becomes a Target Attribute. A Similarity Criterion specifies the requisite degree of similarity. An example is that the mean virtual experiment measurement falls within ± 1 standard deviation of the mean wet-lab measurement. By increasing the strength and variety of analogies between measurements made during experiments on Mouse Analog and corresponding measurements made during experiments on mice (left side of Fig. 1D), the credibility of the Liver’s multilevel model mechanisms increases.

**Figure. 2.**
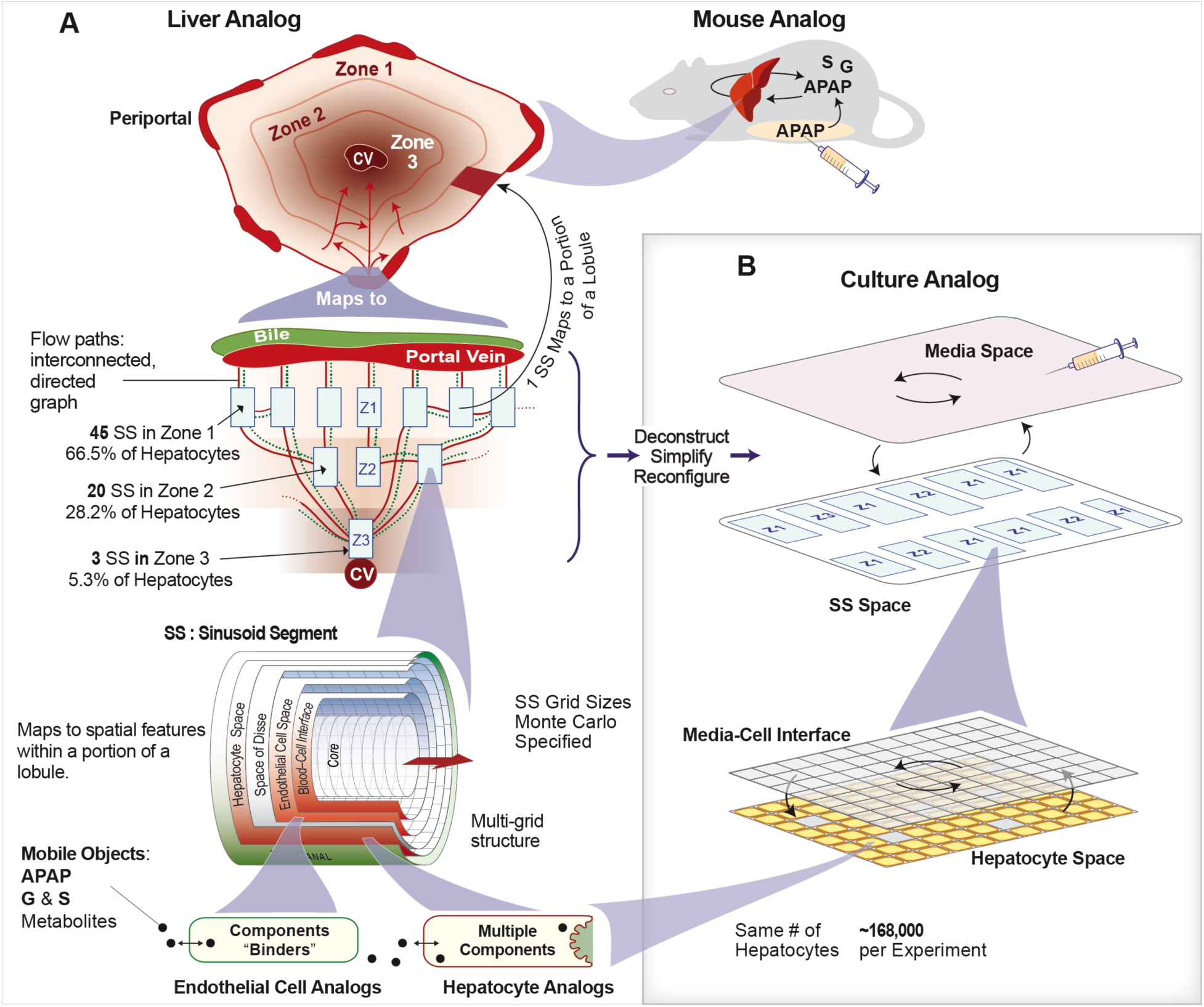
Mouse and Culture Analogs. (A) A Monte Carlo variant of Mouse Analog comprises a nested set of all indicated components. Graph structure (flow paths) and dimensions of Sinusoid Segments within each variant are Monte Carlo-specified. One experiment uses 12 Mouse Analog variants. (B) A Culture Analog variant comprises Media Space and an SS space. All Mouse Lobule SSs are first simplified and then placed in SS space. A simplified SS retains its Hepatocyte Space and Blood-Cell Interface, which is remapped as the Cell-Media Interface. Hepatocyte configurations within Mouse and Culture Analogs are identical. Mobile objects enter and exit spaces stochastically. In Culture, they move between Media and Cell-Media Interface, and between Cell-Media Interface and Hepatocytes.

A Lobule comprises a directed graph with a particular Sinusoid Segment (SS) object (a software agent) at each graph node. SS dimensions are Monte Carlo-determined within constraints that we refined as we achieved additional Target Attributes. Graph nodes are organized into three Zones. Intra-Zone edges within Zones 1 and 2 (there are none in Zone 3) mimic interconnections among sinusoids. Numbers of intra- and inter-Zone edges are fixed, but their node-to-node assignment is Monte Carlo determined for each execution. All flow paths follow the directed graph. One Lobule maps to a tiny random sample of possible lobular flow paths within a whole liver. Bile (dotted green) flows separately from blood (solid red) but is not a factor for this work.

Each SS functions as an analog of a random sample of a portion of a sinusoid plus adjacent tissue. It comprises Core, Bile Space, and four same-size grids: the Blood-Cell Interface (simply Interface hereafter), Endothelial Cell Space, Space of Disse, and Hepatocyte Space. Cell objects occupy most of Endothelial Cell (99%) and Hepatocyte (90%) spaces. APAP, its Metabolites, and some other Solutes are mobile objects. A fraction of APAP in Body (along with other mobile objects) is transferred to Portal Vein each simulation cycle. From there, APAP enters Core and Interface spaces at the upstream end of all Zone 1 SS. They percolate stochastically through accessible spaces influenced by configuration-controlled local flow. APAP that reaches the distal end of Core and Interface spaces are transferred along a connecting edge to another SS. Mobile objects exit Zone 3 SS into Central Vein, where they get moved to Body.

Entry and exit of Solutes from each Endothelial Cell and Hepatocyte is mediated by the cell according to the Solute’s properties. Endothelial Cells contain Binders that bind and release APAP (maps to non-specific binding). Hepatocytes (~14,000 per Lobule) used previously validated event management modules [Petersen et al., 2014], which control material entry and removal along with binding and the object transformations described in Fig. 3. The order of events is (pseudo) randomized each simulation cycle.

**Figure. 3.**
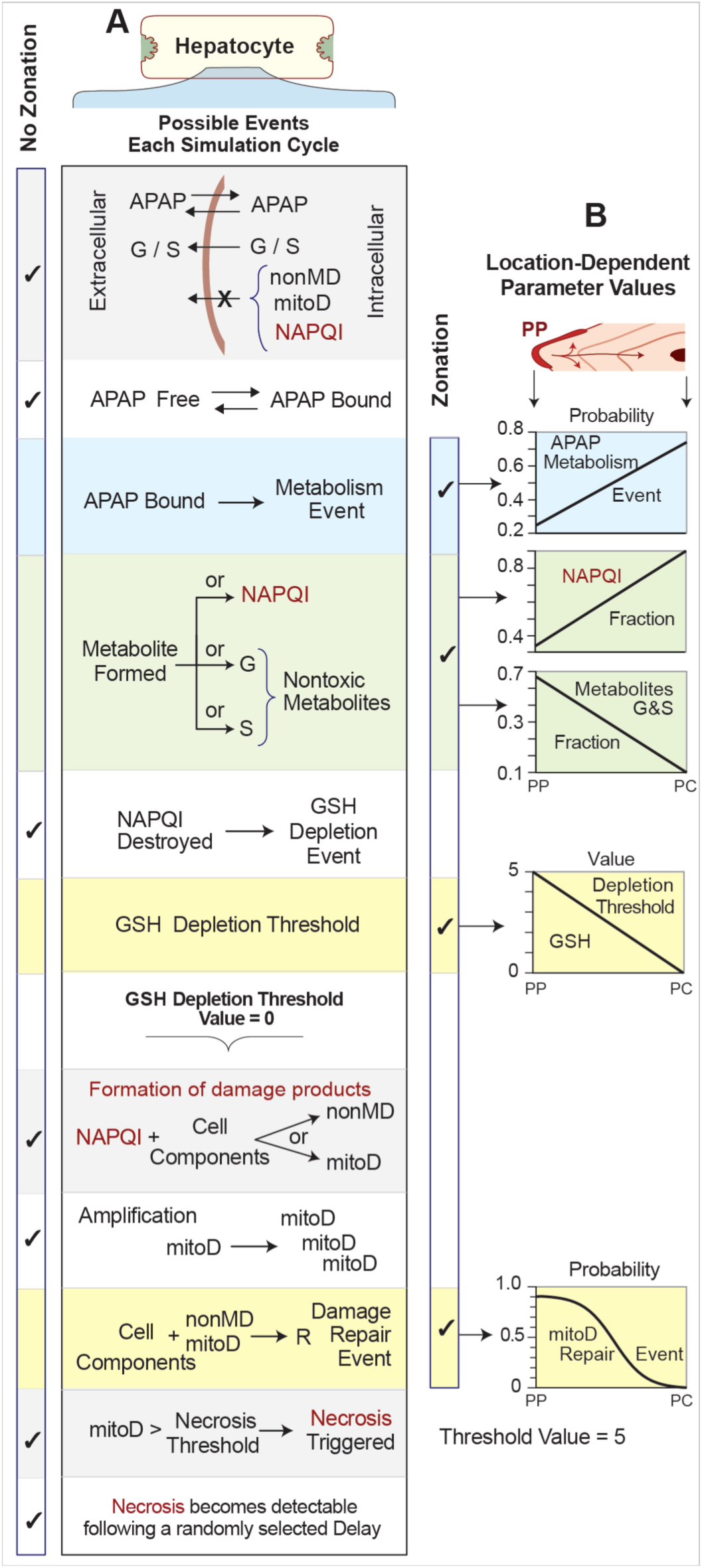
Intra-Hepatocyte events in Mouse and Culture Analogs. (A) Listed under Hepatocyte are the events and activities described in the text. Each event can occur within each Hepatocyte each simulation cycle, as detailed in [Smith et al., 2016]. Each event executes independently in a pseudo-random order. All events are stochastic. Right (Left) side check marks identify events that are (are not) subject to Zonation. When Necrosis is triggered, it becomes measurable (maps to cells that stain positive for necrosis) in the future after some parameter-specified number of simulation cycle. (B) The five graphs show how these events depend on location. PVE = value at Portal Vein entrance; CVE = value at Central Vein exit. First and last graph: probability of event occurrence. Second and third graph: relative metabolite fraction. Fourth graph: value of GSH Depletion threshold.

### Culture Analog

Petersen et al. [Petersen et al., 2016] used a similar Mouse Analog along with 2D models of hepatocyte cultures to explore and challenge mechanism-based hypotheses about immune-mediated P450 down-regulation in vitro. The Hepatocytes utilized in the two systems contained the same internal components. However, because Petersen et al. parameterized them separately, they simulated rather than explain the in vitro-in vivo disconnect (Fig. 1C) within the validation data.

Culture Analog is illustrated in Fig. 2B. It uses Media Space and SS Space. Media Space mimics a well-mixed system. Culture Analog does not use the Lobule’s directed graph. All SSs from a Lobule are simplified, reinstantiated, and then placed into Culture’s SS Space. A Culture Analog does not use Core and Bile spaces, nor does it use Endothelial Cell and Disse grids. It does use Hepatocyte and Interface grids. We remap the latter as the Media-Cell interface. Each simulation cycle, a small fraction of APAP (and other specified Solutes) in Media is transferred randomly to each Media-Cell Interface grid. From there Solutes move within Hepatocyte Space and enter and exit Hepatocytes exactly as they do in Mouse Analog Lobules. Unless specified otherwise, Mouse and Culture Hepatocyte Mechanisms use the same location-specific configuration values (Fig. 3B).

### Intra-Hepatocytes Mechanisms

Objects within Hepatocytes and their capabilities are identical to those used previously [Smith et al., 2016]. The event descriptions that follow are per simulation cycle. Hepatocytes contain four types of physiomimetic modules [Petersen et al., 2014]: InductionHandler (not used in this work), EliminationHandler, MetabolismHandler, and BindingHandler.

An APAP object maps to a tiny fraction of an actual APAP Dose. There is a direct mapping between the probability of an APAP metabolism event and amounts of metabolic enzymes. Both the probability of an APAP metabolic event and the probability that the Metabolite is NAPQI increase threefold from Portal Vein entrance to Central Vein exit. All other metabolites are lumped together and divided equally between G&S (maps to the glucuronide and sulfate metabolites). A Mechanism uses particular values from each of the Fig. 3B gradients (from Portal Vein entrance to Central Vein exit). Each gradient is implemented explicitly as a function of distance from PP entrance to the Hepatocyte’s position. The APAP Liver extraction ratio measured at steady-state (with constant rate APAP infusion) averages 0.5.

With probability to react *=* 0.5, each NAPQI undergoes a reaction. At early times, each NAPQI reaction decrements the Hepatocyte’s GSH Depletion Threshold value by 1.0, which maps to depleting a fraction of a hepatocyte’s available GSH. Each Hepatocyte has a location-determined GSH Depletion Threshold value. A small or zero Threshold value means that Hepatocyte is most sensitive to NAPQI- caused hepatotoxicity. After the GSH Depletion Threshold is breached, NAPQI reacts to form one of two types of Damage products. We parsimoniously specified these Damage products: “mitochondrial damage products,” called mitoD (maps to mitochondrial damage), and “non-mitochondrial damage products,” called nonMD (maps to all other types of damage).

The downstream resolution of events triggered by Damage products is inadequate to simulate toxicity phase events. Mindful of our strong parsimony guideline, we specified that one NAPQI → (1 + *n*) mitoD, where *n* is a pseudo-random draw from the uniform [1, 6]. MitoD amplification also maps to the accumulation of reactive oxygen/nitrogen species. Cell Death (Necrosis) is triggered when the amount mitoD > Necrosis Trigger Threshold value. Once an aHPC designated Dean (necrotic), it stops Metabolizing APAP.

Cell Death always follows a Necrosis Trigger event. Because necrosis is a process, there is a delay between the triggering event and when necrosis becomes detectable in stained tissue sections. Death Delay maps to that process. Thus, the time of a detectable aHPC Death = time of Trigger event + a pseudo-random draw from uniform [1.2, 12] hours.

Hepatocytes utilize multiple mechanisms to mitigate or reverse different types of damage. Consistent with our strong parsimony guideline, we implemented a single mitigation Mechanism, repurposing a Metabolism Module, and named it Repair. It maps to a conflation of all actual mitigation/recovery mechanisms. With a location-specified probability, a Repair object replaces a Damage product. We focused on mitoD because only it can trigger Necrosis. By specifying that the probability of a mitoD Repair event decreases sigmoidally from Portal Vein entrance to Central Vein exit, we enabled Necrosis Trigger events to occur first close to Central Vein, which was the key Targeted Attribute in [Smith et al., 2016].

### Requirements

We use the virtual experiment approach described by Kirschner et al. [Kirschner et al., 2014] along with the enhanced strategies detailed by Petersen et al. [Petersen et al.; 2016, Petersen and Hunt, 2016]. The model mechanisms and methods require meeting the following four requirements.

1. Mouse and Culture Analogs during execution must exhibit all five primary characteristics of a biologically explanatory mechanism [Darden, 2008]: 1) the mechanism is biomimetic and responsible for the phenomena; 2) the mechanism has components: modules, entities, and activities; 3) components are arranged spatially and can exhibit structure, localization, orientation, connectivity, and compartmentalization; 4) activities have temporal aspects, including rate, order, duration, and frequency; and 5) the mechanism has a context, which can include being in a series and/or a hierarchy.
2. Components and spaces (Fig. 2) are concrete, biomimetic [Hunt et al., 2011, Pogson et al., 2006], and sufficiently modular to facilitate analogical reasoning [Bartha, 2013, Frigg and Hartmann, 2012].
3. Phenomena measured at a higher level or layer of organization arise mostly from local component interactions at a lower level of organization.
4. Each mobile object type maps to a particular chemical entity. Quasi-autonomous components (i.e., software agents such as SS and Hepatocytes) recognize different mobile objects and adjust their response appropriately. For example, an Hepatocyte recognizes that an adjacent object has the property *membraneCrossing = yes,* and allows it to enter stochastically.

To achieve Requirement 2, Mouse Analogs are written in Java, utilizing the MASON multi-agent simulation toolkit [Luke et al., 2005].

### Analog technical details

Analogs are treated as a form of data, using both the implicit schema of Java, JavaScript, and R and the explicit schema of the configurations. Mouse Analogs and configuration files are managed using the Subversion version control tool in two repositories, one private (Assembla) and another public. The data presented herein along with Mouse and Culture Analog code are available (https://simtk.org/home/isl/).

The entire toolchain, including the operating system, configurations, and I/O handling is open-source. All project generated released data is available to be licensed as open data. We execute virtual experiments using local hardware and in a cloud environment. Experiments described herein were run using local hardware and virtual machines [Ropella and Hunt, 2010] on Google Compute Engine, running 64-bit Debian 7. Analog quality assurance and control details are discussed in [Smith et al., 2016] along with practices followed for validation, verifications, sensitivity analyses, and uncertainty quantification.

## Results

### Dose-response curves

We use the occurrence of Necrosis Trigger events as our measurement of response. In both Mouse and Culture Analogs, cumulative Trigger events reach a plateau within 180 minutes. We conducted Dose-Response (D-R) experiments (12 Monte Carlo variants). We exposed aHPCs to identical Doses of APAP ranging from 6,000 to 500,000 (largest APAP Dose) objects per Monte Carlo variant. Figure 4 displays results. Dose-dependent differences were also observed (not shown) in measurements of other attributes. Differences in occurrence of Necrosis Trigger events between Mouse and Culture Analog experiments is a complex consequence of differences in aHPC exposure to APAP and aHPC heterogeneity (discussed below). In Culture, the organizational structure of Lobules is absent. Consequently, on average, aHPCs are exposed to APAP that is distributed uniformly within the Media-Cell interface grids. Within a Mouse Lobule, however, aHPCs are exposed to APAP sequentially. During a given interval, upstream and downstream aHPCs are “seeing” different amounts of extra-Hepatocyte APAP. Differences in exposure dynamics between Mouse and Culture Analogs alter intra-Hepatocyte phenomena. The different D-R curves in Fig. 4 are a consequence.

**Figure. 4.**
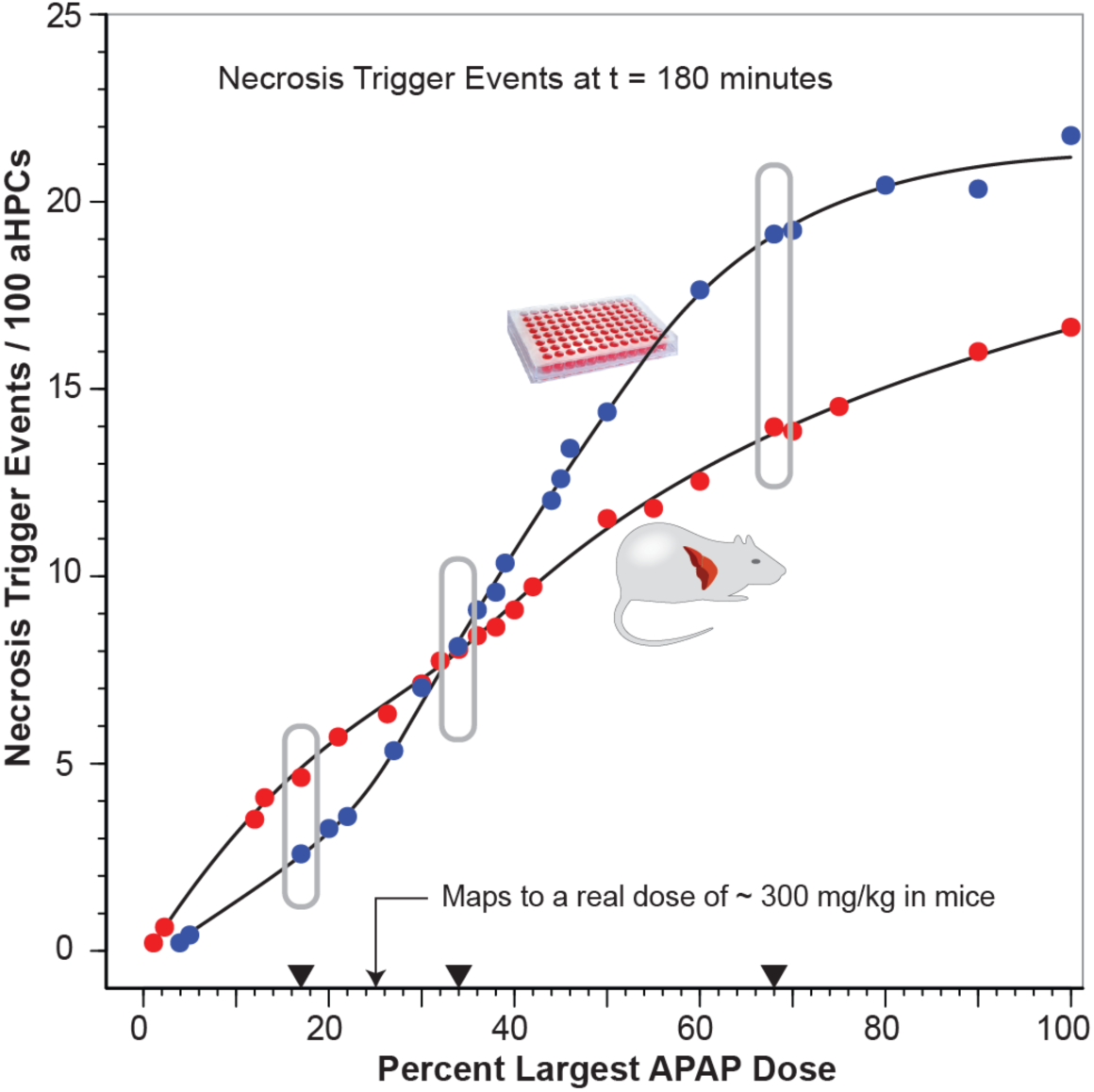
Dose-Response curves for Mouse and Culture Analogs. Dose range: 6,000–500,000 APAP objects per variant. Experiments were conducted separately by two coauthors using different local computers and cloud virtual machines and with different sets of random number seeds. Figures. 5 and 6 present measurements of intra-Hepatocyte phenomena at the three indicated Doses.

### Hepatocyte-level APAP disposition and toxicity measurements

At the end of each simulation cycle (1 second), we measured the number ofNecrosis Trigger events. Virtual measurements are made analogous to wet-lab counterparts to facilitate comparisons when wet-lab data are available. Crossing Dose-response curves indicate that Mechanism entities and activities are contributing differently to cumulative Necrosis Trigger events from smallest to largest Doses. To help identify and explain those differences, measurements of selected phenomena, averaged over all Liver aHPCs, are plotted in Fig. 5. Simulated IP dosing causes peak APAP amounts in the Mouse to occur later than in Culture. For each row of plots in Fig. 5, compare the temporal profile in Mouse to that in Culture for each Dose. Cumulative Necrosis Trigger events in Mouse are greater than in Culture at the smallest Dose, but that relationship is reversed at the largest Dose. For APAP in aHPCs, the Mouse-to-Culture profile relationships are about the same for all three Doses. That is not the case for NAPQI, GSH Depletion events, and Mitochondrial Damage.

**Figure. 5.**
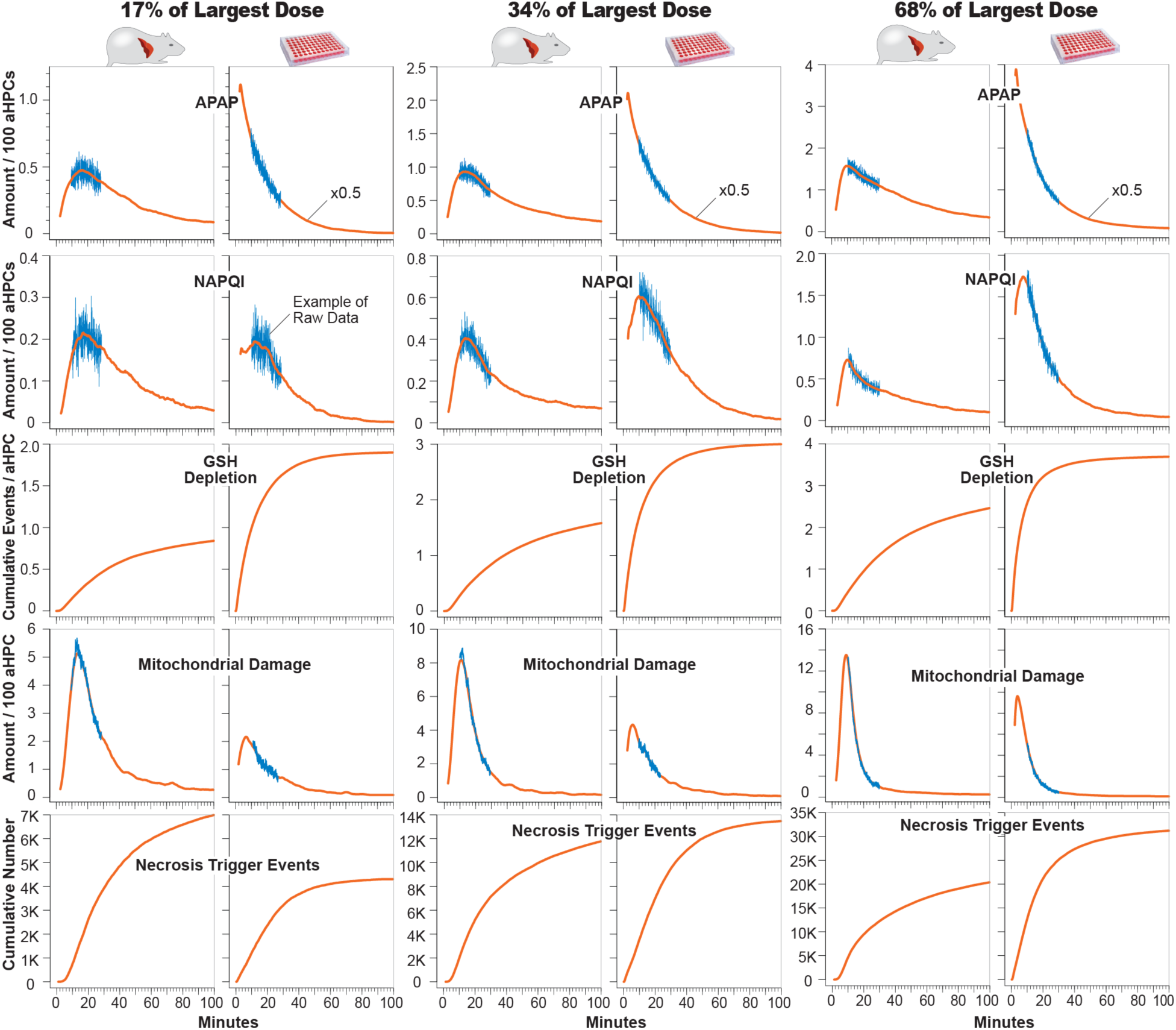
Selected phenomena in Mouse and Culture Analogs. Each row displays temporal profiles for relevant phenomenon measured each second during Mouse and Culture experiments at the three Doses indicated in Fig. 4. The software configuration values for each aHPC are the same for Mouse and Culture. The range of y-axis values increases with increasing Dose. Values for amount/100 aHPCs are 100-point centered moving averages. The actual per second values between 10 and 30 minutes are plotted in blue to illustrate within-experiment variance. Relative variance for smaller-Dose experiments is larger because there are fewer objects and fewer events. The corresponding variance is not evident in cumulative values; however, a repeat experiment would not generate the exact same cumulative value profile. Largest Dose: 500,000 APAP per execution; 12 executions/experiment.

At comparable times, the differences in NAPQI amounts in Culture aHPCs, relative to those in Mouse, increase with increasing APAP Dose. We see a similar trend for Mitochondrial Damage. However, that trend is reversed for GSH Depletion events. At the smallest Dose, relative to results in Culture, fewer Mouse GSH Depletion events caused a larger number of Necrosis Trigger events. That observation seems counterintuitive. At the largest Dose, we do not see that difference: for both Mouse and Culture, the relationship between cumulative GSH Depletion events and cumulative Necrosis Trigger events is similar. Explanations of those results, including the counterintuitive observation, require the more detailed, Lobular-location dependent information provided in Fig. 6.

**Figure. 6.**
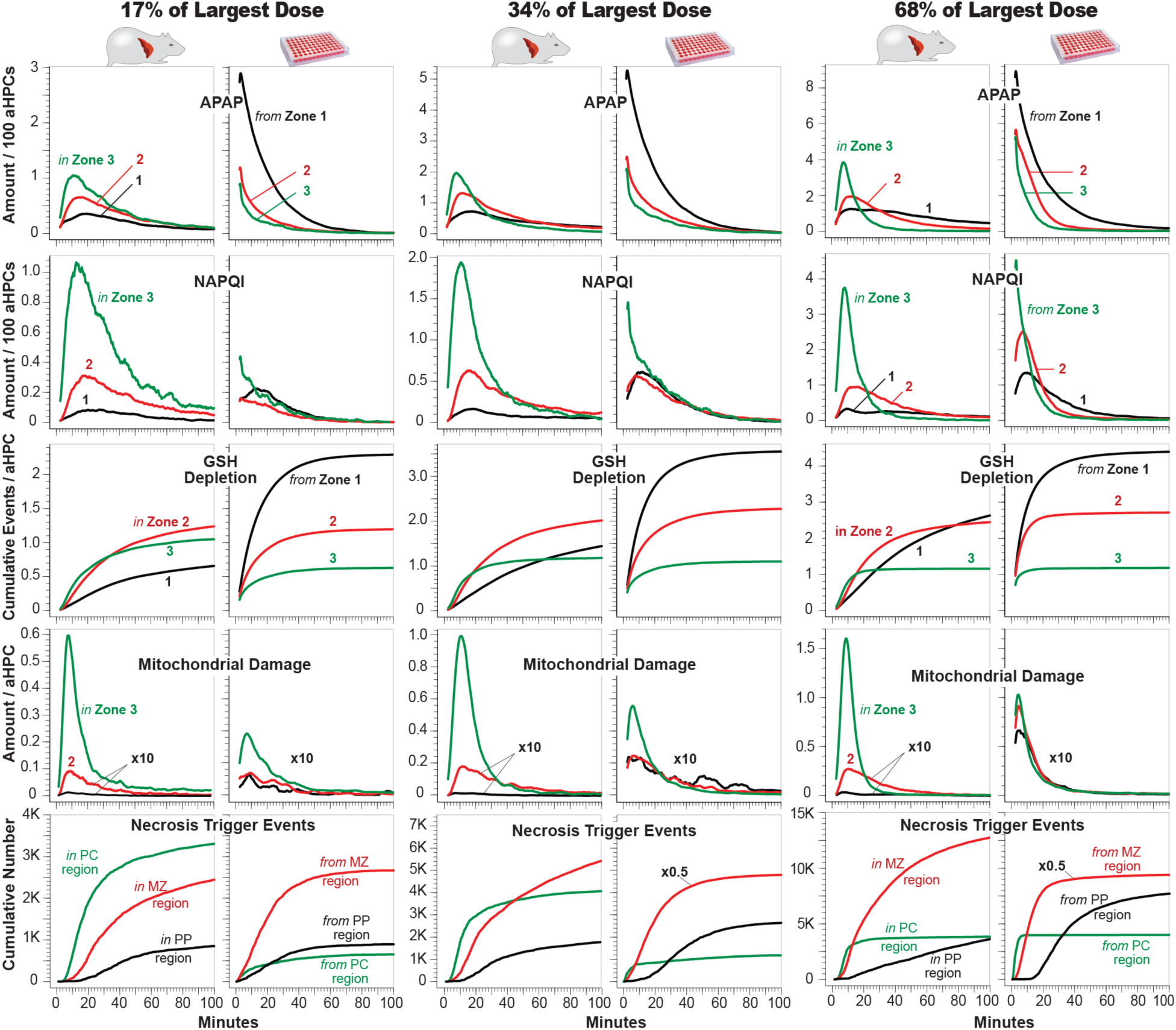
Hepatocyte-level attributes in Mouse and Culture Analogs by Zone. The data are from the same experiments as in Fig. 5. Except for Necrosis Trigger events (see text), measurements are plotted for aHPCs in each SS Zone (e.g., *in* Zone 3, 2 or 1) for Mouse. For Culture, *Zone* refers to the location of those aHPCs within the Lobule before they were “isolated” (e.g., *from* Zone 1, 2 or 3) and reconfigured (Fig. 2B) into Culture Analog. By measuring average values per SS, we can compare how similarly configured aHPCs in Mouse and Culture contexts behave during exposure to APAP. Cumulative number of Trigger events: PC = Pericentral (location is < 10 grid spaces from Central Vein exit); MZ = Midzonal (10 < location is < 20 grid spaces from Central Vein exit); and PP = Periportal (20 < location is < 30 grid spaces from Central Vein exit). The range of y-axis values increases with increasing Dose. Values for amount/100 aHPCS and average Trigger event locations are 100-point centered moving averages.

Amounts of APAP, NAPQI, and Mitochondrial Damage per aHPC, along with GSH Depletion events were measured separately in SSs in Zone 1, 2, and 3. We measured Trigger events in Pericentral, Midzonal, and Periportal regions (defined in the legendto Fig. 6) and plotted them in Fig. 6. For Mouse, values for APAP and GSH Depletion in Zone 3 during the first 10 minutes are largest for all three Doses, whereas, in Culture, the Zone 1 values are largest. That striking difference highlights Mechanism differences between aHPCs in Mouse and Culture and contributes to an explanation of the virtual Mouse-Culture disconnect. That difference is a consequence of two factors: 1) the sequential versus simultaneous aHPC exposure described above. 2) The majority of Lobular aHPCs are in Zone 1. For each Lobule variant, there are 45 SS in Zone 1, 20 in Zone 2, and 3 in Zone 3. That structure maps directly to the polyhedral nature of lobules. Compounds in blood entering portal vein tracts get exposed to many more hepatocytes than blood exiting the central vein. In Liver Analogs, the Zone1/Zone 3 aHPC ratio averages about 12.5, with 66.5% of aHPCs in Zone 1 and 5.3% Zone 3. Consequently, APAP/aHPC in Zone 3 is > than in Zones 2 and 1. The ratio would be larger except for a mitigating factor: as specified in Fig. 3B, the probability of an APAP Metabolism event is Zone 3 > Zone 2 > Zone 1; Zone 3 aHPCs are drained of APAP faster than Zone 1 aHPCs. In Culture, one might expect APAP/aHPC to be about the same because Unbound APAP is directly proportional to local APAP amounts within the Media-Cell Interface grid. However, APAP/aHPC values for Zones 2 and 3 are smaller primarily because the probability of an APAP Metabolism event is larger in Zones 2 and 3 than in Zone 1.

Because NAPQI formation increases periportal-to-pericentral (Fig. 3B), the amounts of NAPQI/aHPC in Mouse are largest in Zone 3. Only at early times, is the same true for Culture. At comparable times after 20 minutes, compare Mouse-to-Culture NAPQI/aHPC at each Dose. For the smallest Dose, the values for Zone 2 and 3 are larger for Mouse. For the largest Dose, the values for Zones 1 and 2 are greater for Culture. That reversal is caused primarily by a combination of two factors: the sequential versus simultaneous aHPC exposure and by 10 minutes, all Zone 3 aHPC in both Mouse and Culture have experienced Necrosis Trigger events.

By 30 minutes after the small APAP Dose, Mouse GSH Depletion events in Zone 2 begin exceeding those in Zone 3. That transition is a consequence of two factors: 1) as plotted in Fig. 3B, average GSH Depletion Threshold values are Zone 2 > Zone 3. So, more GSH Depletion can occur in Zone 2. 2) Depletion events accumulate more slowly in Zone 2 because less APAP got Metabolized to NAPQI in Zone 2 relative to Zone 3. In Culture, cumulative Depletion events are largest for Zone 1 because Depletion Threshold values for Zone 1 aHPCs are largest; there is more GSH to get depleted.

For Mouse and Culture, the relative per Zone profile patterns for the amount of mitoD/aHPC are similar to those for NAPQI except that amount of mitoD/aHPC in Zones 2 and 3 are lower than in Zone 3 by more than 10x. That large difference is because of greater rates of Damage Repair in Zones 1 and 2 limits the accumulation of mitoD.

To see why the D-R curves cross, first, consider the high Dose cumulative Trigger event profiles. Within 30 minutes, Necrosis gets triggered in all Pericentral aHPCs in Mouse and Culture. Because of the more rapid Periportal depletion of Culture GSH relative to Mouse GSH, Trigger events accumulate faster in Culture in Periportal aHPCs. By 60 minutes, Necrosis has also been triggered in all Midzonal aHPCs in Culture, whereas in Mouse, Midzonal aHPCs values are still increasing. The situation is entirely different for cumulative Trigger event profiles following the small Dose. In Mouse, Pericentral aHPCs, which are most sensitive to NAPQI toxicity, experience greater APAP exposures than do upstream aHPCs. That is not the case in Culture. Relative to Mouse, exposure per aHPC is the same in Culture.

For Mouse at all three Doses, early Trigger events occur close to Central Vein exit in the Pericentral region. At t = 100 for the smallest Dose, Midzonal Trigger events in both Mouse and Culture are similar, yet in Mouse, Trigger events are even larger in Pericentral aHPCs, whereas in Culture, they are smallest. Further, Necrosis in Mouse has been Triggered in most Pericentral aHPCs, whereas in Culture, Trigger events have occurred in only a fraction of Pericentral aHPCs. Those differences contribute to an explanation of the virtual Mouse-Culture disconnect.

## Discussion

Software mechanisms become scientifically interesting and useful as analogies when temporal measurements of a generated phenomenon are quantitatively similar to wet-lab counterparts, within some prespecified criterion. Mouse Analog’s Liver and Hepatocytes are analogous to an actual liver and hepatocytes, but only to the extent needed to achieve validation by prespecified Targeted Attributes [Smith et al., 2016]. This same characterization applies to components within aHPCs (Fig. 3) and the multilevel model Mechanisms used to simulate targeted APAP hepatotoxicity phenomena. We embedded Model mechanism knowledge along with structural information within Mouse and Culture analogs. Comparison of virtual experiments on the Mouse and Culture Analogs can help identify and explain mechanistic similarities and differences between in vitro and in vivo systems. Information obtained in these comparisons are expected to enable exploring possible explanations for a known or suspected disconnect and can be used to incrementally develop scientifically challengeable theories (plausible explanations) that may explain the disconnect.

To fully understand the behaviors of hepatocytes in an experimental context, we must take into account various relevant contexts in which they function to generate phenomena of interest, and how function and phenomena change according to context. To advance that understanding, we must consider the methods and processes used to engineer an in vitro (and other) experimental system from whole animal sources. The methods used herein are intended to contribute to that larger effort. The information needed to begin closing or bridging the in vitro–in vivo disconnect cannot be obtained using wet-lab methods alone because complete hepatocyte behavior cannot be measured either in vitro or in vivo, and changes in hepatocyte states during isolation are challenging to control. We suggest that one can begin bridging that disconnect by hypothesizing, instantiating, and then exploring plausible model Mechanisms for Culture and Mouse contexts. The complemental methods presented here use virtual experiments to measure changes that occur when all Hepatocytes from a Mouse Analog’s Liver are precisely translated into a 2D configuration, which is strongly analogous to hepatocytes in 2D cultures. The result is a Culture Analog. We learned about Mouse-Culture disconnects by comparing the similarities and differences in how the two systems respond to simulated Doses of APAP.

Bridging wet-lab disconnects will be considerably more challenging than bridging a virtual disconnect, so we first consider how the latter might be accomplished. Given only the Culture data in Figs. 4 and 5, but no details about Culture Mechanism, we face a virtual in vitro-in vivo disconnect: it is infeasible to predict (within some reasonable tolerance) cumulative Necross Trigger events within Mouse Analogs, even though Culture data in Fig. 5 is quite detailed. Even knowing that experiments in Figs. 4 and 5 met requisite four in the Introduction (i.e., same intra-Hepatocyte mechanisms), we need some measure of aHPC homogeneity, or lack thereof, in Culture to begin constructing a Mechanism-based theory to bridge across the disconnect. An example of a virtual measurement of aHPC heterogeneity: create many non-membrane-crossing mobile Reporter objects. Include them with the APAP Dose. When adjacent to an aHPC, a Reporter object asks the aHPC, is the value of your GSH Depletion Threshold ≥ 2? If no, the Reporter takes on a value of 0 and then departs; if yes, it asks, is your Threshold ≥ 4? If no, it takes on a value of 1; if yes, it takes on a value of 2 and leaves. Each aHPC only responds once to a Reporter object. Given the cumulative fraction of Dosed Reporters having values 0, 1, and 2, we can estimate the relative numbers of pericentral, midzonal, and pericentral aHPCs in that Culture.

It is feasible to obtain similar wet-lab information, e.g., by measuring the percent of cells that stain positive for a periportal or pericentral location marker. In general, results from wet-lab measurements that provide information about the heterogeneity of relevant mechanism component parts, component operations, and their organization would be among the “multi-attribute similarities” requested on the right side of Fig. 1D. However, there are no wet-lab measurements that will recover lost lobular structural and organizational properties that influence the phenomena of interest in vivo. Lobular architecture and intra- aHPC mechanisms are entangled within lobules and cannot be separated. Having achieved multi-attribute similarities between Mouse and Liver Analogs and wet-lab measurements for multiple compounds, we use the ability to transition smoothly between Culture and Mouse Analogs to simulate that entanglement.

Once we establish quantitative multi-attribute similarities between Culture Analog and wet-lab counterparts, we can reconfigure Culture Analog aHPCs into an organized Lobule structure within Mouse Analog. The intra-aHPC model mechanisms will be the same. Virtual experiments on the resulting Virtual Mouse Analogs can provide plausible model mechanism-based predictions of corresponding experiments in mice. The credibility of those predictions will be wholly dependent on the credibility of the model mechanisms and multi-attribute similarities established on the right and left sides of Fig. 1D. The virtual Culture-to-Mouse translation can stand as a concrete, scientifically challengeable theory explaining the in vitro-in vivo disconnect. That model mechanism-based theory provides the framework to develop more reliable interpretations of in vitro observations, which then may be used to improve extrapolations.

In vitro to in vivo extrapolation is a method to bridge the in vitro-in vivo disconnect. Such extrapolation is similar to the translation discussed above: mapping measurements from one model to another using assumptions above missing or unknown information (spatial, structural, networking, mechanism gradient, etc.). For some xenobiotics, this information is available or can be approximated using virtual experiments. In those cases, we can generate sufficiently reliable extrapolations (predictions). However, for other xenobiotics, it may not be feasible. By tightly coupling virtual and wet-lab methods we can identify characteristic xenobiotic traits that distinguish the two sets.

## Acknowledgements

The UCSF BioSystems group supported this work. Authors thank Ryan Kennedy for constructive criticism.

## Author Contributions

Conceived and designed the experiments: Smith, Ropella, Hunt

Performed the experiments: Smith, Xu

Analyzed the data: Smith, Xu

Wrote the paper: Smith, Hunt, Ropella

Developed and verified Analog and sub-component requirements: Ropella, Smith

Guided refinements of analog to mouse and culture mappings: Smith, Ropella

Designed and executed batch sampling: Smith, Ropella, Xu

Managed and executed cloud experiments: Smith, Xu Managed current versioning: Ropella

Conducted unit tests: Smith, Ropella

Developed scripts for data analyses and data visualizations: Xu, Smith, Ropella

Conducted component/module validation experiments: Smith

Contributed manuscript content: Smith, Xu, Hunt

Produced figures: Hunt

## References

Bartha P. Analogy and Analogical Reasoning. The Stanford Encyclopedia of Philosophy (Fall 2013 Edition). 2013. Available: http://plato.stanford.edu/archives/fall2013/entries/reasoning-analogy/

Bowman CM and Benet LZ (2016) Hepatic Clearance Predictions from In Vitro-In Vivo Extrapolation and the Biopharmaceutics Drug Disposition Classification System. Drug Metabolism and Disposition 44: 1731–1735.

Darden L (2008) Thinking again about biological mechanisms. Philosophy of Science 75: 958–69.

Fraczek J, Bolleyn J, Vanhaecke T, Rogiers V, and Vinken M (2013) Primary hepatocyte cultures for pharmaco-toxicological studies: at the busy crossroad of various anti-dedifferentiation strategies. Archives of Toxicology 87: 577–610.

Frigg R, Hartmann, S. Models in Science. The Stanford Encyclopedia of Philosophy (Fall 2012 Edition). 2012. Available: http://plato.stanford.edu/archives/fall2012/entries/models-science/.

Godoy P, Hewitt NJ, Albrecht U, Andersen ME, Ansari N, Bhattacharya S, Bode JG, Bolleyn J, Borner C, Böttger J, Braeuning A, Budinsky RA, Burkhardt B, Cameron NR, Camussi G, Cho CS, Choi YJ, Craig Rowlands J, Dahmen U, Damm G, Dirsch O, Donato MT, Dong J, Dooley S, Drasdo D, Eakins R, Ferreira KS, Fonsato V, Fraczek J, Gebhardt R, Gibson A, Glanemann M, Goldring CE, Gómez-Lechón MJ, Groothuis GM, Gustavsson L, Guyot C, Hallifax D, Hammad S, Hayward A, Häussinger D, Hellerbrand C, Hewitt P, Hoehme S, Holzhütter HG, Houston JB, Hrach J, Ito K, Jaeschke H, Keitel V, Kelm JM, Kevin Park B, Kordes C, Kullak-Ublick GA, LeCluyse EL, Lu P, Luebke-Wheeler J, Lutz A, Maltman DJ, Matz-Soja M, McMullen P, Merfort I, Messner S, Meyer C, Mwinyi J, Naisbitt DJ, Nussler AK, Olinga P, Pampaloni F, Pi J, Pluta L, Przyborski SA, Ramachandran A, Rogiers V, Rowe C, Schelcher C, Schmich K, Schwarz M, Singh B, Stelzer EH, Stieger B, Stöber R, Sugiyama Y, Tetta C, Thasler WE, Vanhaecke T, Vinken M, Weiss TS, Widera A, Woods CG, Xu JJ, Yarborough KM, and Hengstler JG (2013) Recent advances in 2D and 3D in vitro systems using primary hepatocytes, alternative hepatocyte sources and non-parenchymal liver cells and their use in investigating mechanisms of hepatotoxicity, cell signaling and ADME. Archives of Toxicology 87:1315–530.

Hunt CA, Ropella GE, Lam TN, and Gewitz AD (2011) Relational grounding facilitates development of scientifically useful multiscale models. Theoretical Biology and Medical Modelling 8: 35–66.

Kirschner DE, Hunt CA, Marino S, Fallahi-Sichani M, and Linderman JJ (2014) Tuneable resolution as a systems biology approach for multi-scale, multi-compartment computational models. Wiley Interdisciplinary Reviews: Systems Biology and Medicine 6: 289–309.

LeCluyse EL, Witek RP, Andersen ME, and Powers MJ (2012) Organotypic liver culture models: meeting current challenges in toxicity testing. Critical Reviews in Toxicology 42: 501–48.

Luke S, Cioffi-Revilla C, Panait L, Sullivan K, and Balan G (2005) MASON: A Multiagent Simulation Environment. Simulation 81: 517–27.

Park S, Ropella GEP, Kim SH, Roberts MS, and Hunt CA (2009) Computational strategies unravel and trace how liver disease changes hepatic drug disposition. Journal of Pharmacology and Experimental Therapeutics 328: 294–305.

Park S, Kim SH, Ropella GEP, Roberts MS, and Hunt CA (2010) Tracing multiscale mechanisms of drug disposition in normal and diseased livers. Journal of Pharmacology and Experimental Therapeutics 334: 124–136.

Petersen BK, Ropella GE, and Hunt CA (2014) Toward modular biological models: defining analog modules based on referent physiological mechanisms. BMC Systems Biology 8: 95–112.

Petersen BK, Ropella GE, and Hunt CA. (2016) Virtual experiments enable exploring and challenging explanatory mechanisms of immune-mediated P450 down-regulation. PloS ONE 11: e0155855.

Petersen BK and Hunt CA. (2016) Developing a vision for executing scientifically useful virtual biomedical experiments. in Proceedings of the 2016 Spring Simulation Multiconference. p. 697–706, Society for Computer Simulation International, San Diego.

Pogson M, Smallwood R, Qwarnstrom E, and Holcombe M (2006) Formal agent-based modelling of intracellular chemical interactions. Biosystems 85: 37–45.

Poulin P (2016) The Need for Human Exposure Projection in the Interpretation of Preclinical In Vitro and In Vivo ADME Tox Data, in Drug Discovery Toxicology: From Target Assessment to Translational Biomarkers (eds Y. Will, J. E. McDuffie, A. J. Olaharski and B. D. Jeffy), John Wiley & Sons, Inc, Hoboken, NJ.

Ropella GEP and Hunt CA (2010) Cloud computing and validation of expandable In silico livers. BMC Systems Biology 4: 178.

Smith AK, Petersen BK, Ropella GE, Kennedy RC, Kaplowitz N, Ookhtens M, and Hunt CA (2016) Competing Mechanistic Hypotheses of Acetaminophen-Induced Hepatotoxicity Challenged by Virtual Experiments. PloS computational biology 12: e1005253.

Smith AK, Ropella GE, Kaplowitz N, Ookhtens M, and Hunt CA (2014) Mechanistic Agent-based Damage and Repair Models as Hypotheses for Patterns of Necrosis Caused by Drug Induced Liver Injury. in Proceedings of the 2014 Summer Computer Simulation Multiconference p. 112–120, Society for Computer Simulation International, San Diego.

Soldatow VY, LeCluyse EL, Griffith LG, and Rusyn I (2013) In vitro models for liver toxicity testing. Toxicology Research 2: 23–39.

Tetsuka K, Ohbuchi M, and Tabata K (2017) Recent Progress in Hepatocyte Culture Models and Their Application to the Assessment of Drug Metabolism, Transport, and Toxicity in Drug Discovery: The Value of Tissue Engineering for the Successful Development of a Microphysiological System. Journal of Pharmaceutical Sciences 106: 2302–2311.

Vellonen KS, Malinen M, Mannermaa E, Subrizi A, Toropainen E, Lou YR, Kidron H, Yliperttula M, Urtti A (2014) A critical assessment of in vitro tissue models for ADME and drug delivery. Journal of Controlled Release 190: 94–114.

Yan L, Sheihk-Bahaei S, Park S, Ropella GE, and Hunt CA (2008a) Predictions of hepatic disposition properties using a mechanistically realistic, physiologically based model. Drug Metabolism and Disposition 36: 759–768.

Yan L, Ropella GE, Park S, Roberts MS, and Hunt CA (2008b) Modeling and simulation of hepatic drug disposition using a physiologically based, multi-agent in silico liver. Pharmaceutical Research 25: 1023–1036.

